# Refining the accuracy of validated target identification through coding variant fine-mapping in type 2 diabetes

**DOI:** 10.1101/144410

**Authors:** Anubha Mahajan, Jennifer Wessel, Sara M Willems, Wei Zhao, Neil R Robertson, Audrey Y Chu, Wei Gan, Hidetoshi Kitajima, Daniel Taliun, N William Rayner, Xiuqing Guo, Yingchang Lu, Man Li, Richard A Jensen, Yao Hu, Shaofeng Huo, Kurt K Lohman, Weihua Zhang, James P Cook, Bram Prins, Jason Flannick, Niels Grarup, Vassily Vladimirovich Trubetskoy, Jasmina Kravic, Young Jin Kim, Denis V Rybin, Hanieh Yaghootkar, Martina Mñller-Nurasyid, Karina Meidtner, Ruifang Li-Gao, Tibor V Varga, Jonathan Marten, Jin Li, Albert Vernon Smith, Ping An, Symen Ligthart, Stefan Gustafsson, Giovanni Malerba, Ayse Demirkan, Juan Fernandez Tajes, Valgerdur Steinthorsdottir, Matthias Wuttke, Cécile Lecoeur, Michael Preuss, Lawrence F Bielak, Marielisa Graff, Heather M Highland, Anne E Justice, Dajiang J Liu, Eirini Marouli, Gina Marie Peloso, Helen R Warren, ExomeBP Consortium, MAGIC Consortium, GIANT Consortium, Saima Afaq, Shoaib Afzal, Emma Ahlqvist, Peter Almgren, Najaf Amin, Lia B Bang, Alain G Bertoni, Cristina Bombieri, Jette Bork-Jensen, Ivan Brandslund, Jennifer A Brody, Noël P Burtt, Mickaël Canouil, Yii-Der Ida Chen, Yoon Shin Cho, Cramer Christensen, Sophie V Eastwood, Kai-Uwe Eckardt, Krista Fischer, Giovanni Gambaro, Vilmantas Giedraitis, Megan L Grove, Hugoline G de Haan, Sophie Hackinger, Yang Hai, Sohee Han, Anne Tybjærg-Hansen, Marie-France Hivert, Bo Isomaa, Susanne Jäger, Marit E Jørgensen, Torben Jørgensen, Annemari Käräjämäki, Bong-Jo Kim, Sung Soo Kim, Heikki A Koistinen, Peter Kovacs, Jennifer Kriebel, Florian Kronenberg, Kristi Läll, Leslie A Lange, Jung-Jin Lee, Benjamin Lehne, Huaixing Li, Keng-Hung Lin, Allan Linneberg, Ching-Ti Liu, Jun Liu, Marie Loh, Reedik Mägi, Vasiliki Mamakou, Roberta McKean-Cowdin, Girish Nadkarni, Matt Neville, Sune F Nielsen, Ioanna Ntalla, Patricia A Peyser, Wolfgang Rathmann, Kenneth Rice, Stephen S Rich, Line Rode, Olov Rolandsson, Sebastian Schönherr, Elizabeth Selvin, Kerrin S Small, Alena Stančáková, Praveen Surendran, Kent D Taylor, Tanya M Teslovich, Barbara Thorand, Gudmar Thorleifsson, Adrienne Tin, Anke Tönjes, Anette Varbo, Daniel R Witte, Andrew R Wood, Pranav Yajnik, Jie Yao, Loïc Yengo, Robin Young, Philippe Amouyel, Heiner Boeing, Eric Boerwinkle, Erwin P Bottinger, Rajiv Chowdhury, Francis S Collins, George Dedoussis, Abbas Dehghan, Panos Deloukas, Marco M Ferrario, Jean Ferrières, Jose C Florez, Philippe Frossard, Vilmundur Gudnason, Tamara B Harris, Susan R Heckbert, Joanna M M Howson, Martin Ingelsson, Sekar Kathiresan, Frank Kee, Johanna Kuusisto, Claudia Langenberg, Lenore J Launer, Cecilia M Lindgren, Satu Männistö, Thomas Meitinger, Olle Melander, Karen L Mohlke, Marie Moitry, Andrew D Morris, Alison D Murray, Renée de Mutsert, Marju Orho-Melander, Katharine R Owen, Markus Perola, Annette Peters, Michael A Province, Asif Rasheed, Paul M Ridker, Fernando Rivadineira, Frits R Rosendaal, Anders H Rosengren, Veikko Salomaa, Wayne H-H Sheu, Rob Sladek, Blair H Smith, Konstantin Strauch, André G Uitterlinden, Rohit Varma, Cristen J Willer, Matthias Blüher, Adam S Butterworth, John Campbell Chambers, Daniel I Chasman, John Danesh, Cornelia van Duijn, Josee Dupuis, Oscar H Franco, Paul W Franks, Philippe Froguel, Harald Grallert, Leif Groop, Bok-Ghee Han, Torben Hansen, Andrew T Hattersley, Caroline Hayward, Erik Ingelsson, Sharon LR Kardia, Fredrik Karpe, Jaspal Singh Kooner, Anna Köttgen, Kari Kuulasmaa, Markku Laakso, Xu Lin, Lars Lind, Yongmei Liu, Ruth J F Loos, Jonathan Marchini, Andres Metspalu, Dennis Mook-Kanamori, Børge G Nordestgaard, Colin N A Palmer, James S Pankow, Oluf Pedersen, Bruce M Psaty, Rainer Rauramaa, Naveed Sattar, Matthias B Schulze, Nicole Soranzo, Timothy D Spector, Kari Stefansson, Michael Stumvoll, Unnur Thorsteinsdottir, Tiinamaija Tuomi, Jaakko Tuomilehto, Nicholas J Wareham, James G Wilson, Eleftheria Zeggini, Robert A Scott, Inês Barroso, Timothy M Frayling, Mark O Goodarzi, James B Meigs, Michael Boehnke, Danish Saleheen, Andrew P Morris, Jerome I Rotter, Mark I McCarthy

## Abstract

Identification of coding variant associations for complex diseases offers a direct route to biological insight, but is dependent on appropriate inference concerning the causal impact of those variants on disease risk. We aggregated coding variant data for 81,412 type 2 diabetes (T2D) cases and 370,832 controls of diverse ancestry, identifying 40 distinct coding variant association signals (at 38 loci) reaching significance (*p*<2.2×10^−7^). Of these, 16 represent novel associations mapping outside known genome-wide association study (GWAS) signals. We make two important observations. First, despite a threefold increase in sample size over previous efforts, only five of the 40 signals are driven by variants with minor allele frequency <5%, and we find no evidence for low-frequency variants with allelic odds ratio >1.29. Second, we used GWAS data from 50,160 T2D cases and 465,272 controls of European ancestry to fine-map these associated coding variants in their regional context, with and without additional weighting to account for the global enrichment of complex trait association signals in coding exons. At the 37 signals for which we attempted fine-mapping, we demonstrate convincing support (posterior probability >80% under the “annotation-weighted” model) that coding variants are causal for the association at 16 (including novel signals involving *POC5* p.His36Arg, *ANKH* p.Arg187Gln, *WSCD2* p.Thr113Ile, *PLCB3* p.Ser778Leu, and *PNPLA3* p.Ile148Met). However, at 13 of the 37 loci, the associated coding variants represent “false leads” and naïve analysis could have led to an erroneous inference regarding the effector transcript mediating the signal. Accurate identification of validated targets is dependent on correct specification of the contribution of coding and non-coding mediated mechanisms at associated loci.

---

Genome-wide association studies (GWAS) have identified many thousands of association signals influencing common, complex traits such as type 2 diabetes (T2D) and obesity^1-7^. Most of these significant association signals involve common variants that map to non-coding sequence and identification of their cognate effector transcripts has often proved challenging. The identification of coding variants causally implicated in trait predisposition offers a more direct route from association signal to biological inference.

The exome occupies only 1.5% of overall genome sequence, but modelling of complex trait architecture indicates that, for many common diseases, coding variants make a disproportionately large contribution to trait heritability^8,9^. This enrichment indicates that coding variant association signals have an enhanced probability of being causal when compared to those involving an otherwise equivalent non-coding variant. This does not, however, guarantee that all coding variant associations are causal. Alleles driving common-variant (minor allele frequency [MAF] ≥5%) GWAS signals typically reside on extended risk haplotypes that, due to linkage disequilibrium (LD), incorporate many common variants^10,11^. Consequently, the presence of a coding allele on the risk haplotype does not constitute sufficient evidence that it represents the causal variant at the locus, or that the gene within which it lies is mediating the association signal. Since much coding variant discovery has proceeded through exome-specific analyses via exome-array genotyping or exome sequencing, researchers have often been poorly-placed to position coding variant associations in the context of regional genetic variation, and it is unclear how often this may lead to incorrect assumptions regarding their causal role.

In our recent study of T2D predisposition^12^, we surveyed the exomes of 34,809 T2D cases and 57,985 controls, of predominantly (>90%) European descent, and identified 13 distinct coding variant associations reaching genome-wide significance. Twelve of these associations involved common variants, but the data hinted at a substantial pool of lower-frequency coding variants of moderate impact (allelic odds ratio [OR] between 1.10 and 2.66) that might be amenable to detection in larger samples. We also reported that, whilst many of these signals fell within common variant loci previously identified by GWAS, it was often far from trivial to determine, using available data, whether those coding variants were causal or simply ‘hitchhiking’ on risk haplotypes.

Here, we report analyses that address these two key issues. First, we extended the scope of our exome-array genotyping study to include data from 81,412 T2D cases and 370,832 controls of diverse ancestry, substantially expanding our power to detect coding variant associations across the allele-frequency spectrum. Second, to understand the extent to which the identification of coding variant associations provides a reliable guide to causal mechanisms, we undertook high-resolution fine-mapping of those coding variant association signals amenable to fine-mapping in European samples, in 50,160 T2D cases and 465,272 controls of European ancestry with genome-wide genotyping data.

## RESULTS

### Discovery study overview

First, we set out to discover coding variant association signals by aggregating T2D association summary statistics in up to 452,244 individuals (effective sample size 228,825) across five ancestry groups, performing both European-specific (EUR, 60.9% of total effective sample size) and trans-ethnic (TE) meta-analyses (**Supplementary Tables 1 and 2**). This discovery analysis was restricted to the 247,470 variants represented on the exome array. Exome array genotypes were gathered from: (a) 58,425 cases and 188,032 controls genotyped with the exome array; (b) 14,608 cases and 174,322 controls from UK Biobank and GERA (Resource for Genetic Epidemiology on Adult Health and Aging) genotyped with GWAS arrays enriched for exome content and/or coverage of low-frequency variation across ethnic groups^13,14^; and (c) 8,379 cases and 8,478 controls with whole-exome sequence from the GoT2D/T2D-GENES^12^ and SIGMA^15^ studies. Overall, this represented a 3-fold increase in effective sample size over our previous study of T2D predisposition within coding sequence^12^. Variation in body mass index (BMI) is an established contributor to the T2D-risk: therefore, association analyses were conducted with and without BMI adjustment (adjBMI) to deconvolute the impact of obesity on T2D-associated variants.

We considered *p*<2.2×10^−7^ as significant for protein truncating variants (PTVs) and moderate impact coding variants (including missense, in-frame indel and splice region variants) based on a weighted Bonferroni correction that accounts for the observed enrichment in complex trait association signals mapping to coding variation^16^. This threshold is close to that obtained through other approaches such as simple Bonferroni correction for the total number of coding variants on the array (**Methods**). Compared to our previous study^12^, the expanded sample size substantially increased power to detect association for common variants of modest effect (e.g. from 14.4% to 97.9% for a variant with 20% MAF and OR=1.05) and lower-frequency variants with larger effects (e.g. from 11.8% to 97.5% for a variant with 1% MAF and OR=1.20) assuming homogenous allelic effects across ancestry groups (**Methods**).

### Insights into coding variant association signals underlying T2D susceptibility

We detected significant associations at 69 coding variants under an additive genetic model (either in BMI unadjusted or adjusted analysis), mapping to 38 loci (**Supplementary Fig. 1, Supplementary Table 3**). We observed minimal evidence of heterogeneity in allelic OR between ancestry groups (**Supplementary Table 3**), and no compelling evidence for non-additive allelic effects, irrespective of allele frequency (**Supplementary Fig. 2, Supplementary Table 4**).

Reciprocal conditional analyses (**Methods**) indicated that the 69 coding variants represented 40 distinct association signals (conditional *p*<2.2×10^−7^) across the 38 loci, with two distinct signals each at *HNF1A* and *RREB1* (**Supplementary Table 5**). These 40 signals included the 13 associations reported in our earlier publication^12^, all of which demonstrated more significant associations in this expanded trans-ethnic meta-analysis (**Supplementary Table 6**). Twenty five of the 40 signals were significant (*p*<2.2×10^−7^) in both EUR and TE analyses. Of the other 15 loci, three (*PLCB3*, *C17orf58*, and *ZHX3*) were significant in EUR alone: all reached *p*<6.8×10^−6^ in the combined TE, and for *PLCB3* and *ZHX3*, risk allele frequencies were substantially lower outside Europeans. Twelve loci (**Supplementary Table 3**) were significant in TE alone, but for these (with the exception of *PAX4* which is East Asian specific), the evidence for association was proportionate in the smaller European component (***P***_EUR_<8.4×10^−5^).

Sixteen of the 40 distinct association signals mapped outside regions previously implicated in T2D susceptibility, defined as >500kb from the reported lead GWAS SNPs (Methods, **Table 1**). These included signals involving missense variants in *POC5* (p.His36Arg, rs2307111, *P*_TE_=1.6×10^−15^), *PNPLA3* (p.Ile148Met, rs738409, ***p***_TEadjBMI_=2.8×10^−11^), and *ZZEF1* (p.Ile2014Val, rs781831, ***P***_TE_=8.3×10^−11^).

**Table 1.**
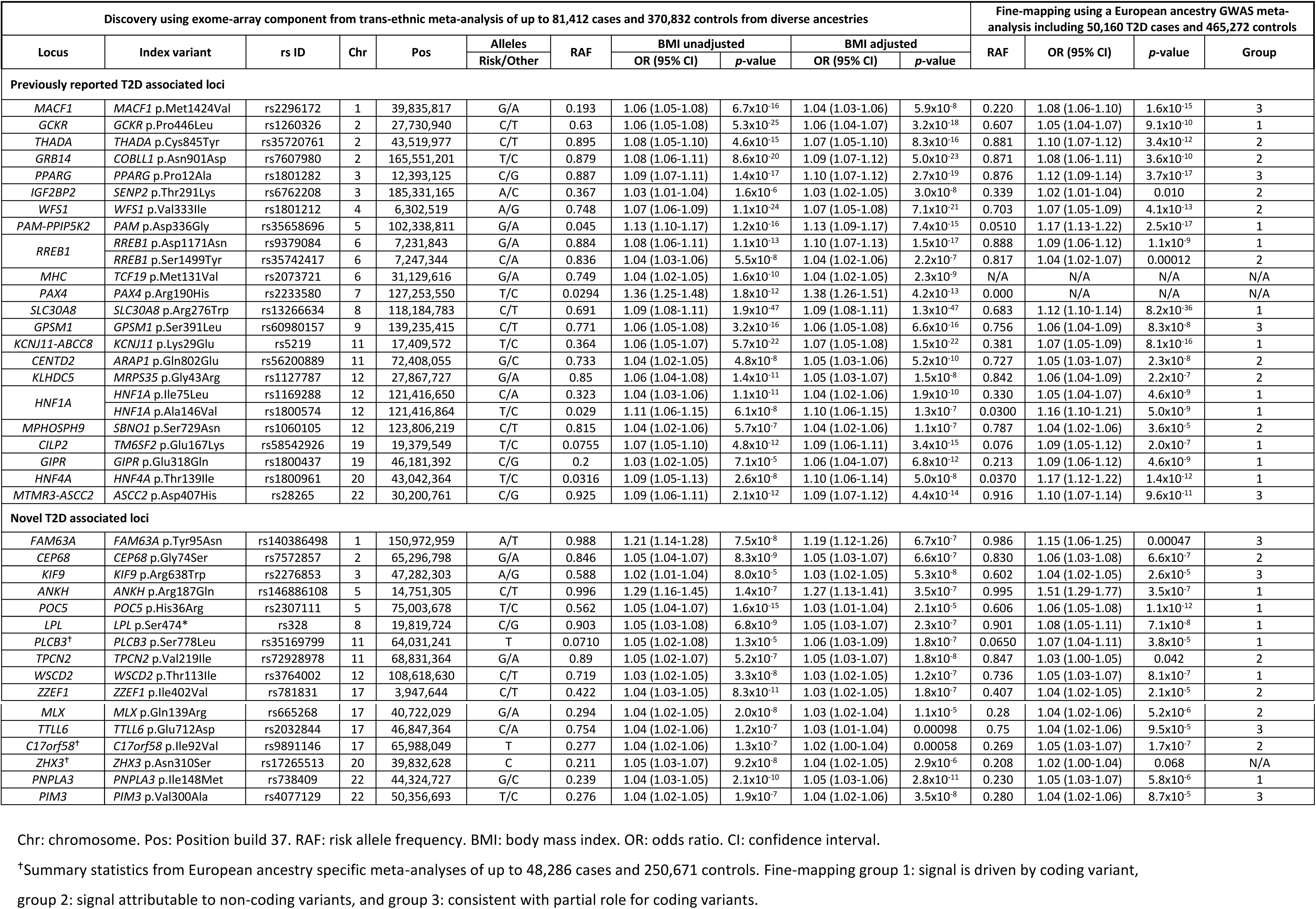
Summary of discovery and fine-mapping analyses of the 40 index coding variants associated with T2D (*p*<2.2×10^−7^).

### Contribution of low-frequency and rare coding variation to T2D susceptibility

Despite increased power and good coverage of low-frequency variants on the exome array (>80% of coding variants with MAF >0.5% in European ancestry populations, as estimated in Fuchsberger et al.^12^), all but five of the 40 distinct coding variant association signals were common, with modest effects (allelic OR 1.02-1.36) (**Supplementary Fig. 3, Supplementary Table 3**). The five association signals attributable to lower-frequency variants were also of modest effect (allelic OR 1.09-1.29) (**Supplementary Fig. 3**). Two of these lower-frequency variant signals were novel, and in both, the minor allele was protective against T2D: *FAM63A* p.Tyr95Asn (rs140386498, MAF=1.2%, OR= 0.82 [0.77-0.88], ***P***_EUR_=5.8×10^−8^) and *ANKH* p.Arg187Gln (rs146886108, MAF=0.4%, OR=0.78 [0.69-87], ***P***_EUR_=2.0×10^−7^). Both variants were very rare or monomorphic in non-European individuals analysed in this study.

In Fuchsberger et al.^12^, we had highlighted a set of 100 low-frequency coding variants with allelic ORs between 1.10 and 2.66, which, despite relatively large estimates for liability-scale variance explained, had not reached overall significance. In this expanded analysis, only five of these variants, including the two novel associations at *FAM63A* p.Tyr95Asn and *ANKH* p.Arg187Gln, achieved significance. More precise effect size estimation with the larger sample size indicates that the OR estimates in the earlier study were subject to a substantial upwards bias (**Supplementary Fig. 3**).

To detect additional rare variant association signals, we performed gene-based analyses (burden and SKAT^17^) using previously-defined “strict” and “broad” masks, filtered for annotation and MAF^18^ (**Methods**). We identified gene-based associations with T2D susceptibility (*p*<2.5×10^−6^, Bonferroni correction for 20,000 genes) for *FAM63A* (SKAT broad mask, 10 variants, combined MAF=1.90%, ***P****_EUR_*=3.1×10^−9^) and *PAM* (SKAT broad mask, 17 variants, combined MAF=4.67%, ***P****_TE_*=8.2x10^-9^). On conditional analysis (**Supplementary Table 7**), we found that the gene-based signal at *FAM63A* was entirely accounted for by the low-frequency p.Tyr95Asn allele described earlier (SKAT broad mask, conditional *p*=0.26). In these data, the gene-based signal for *PAM* is also attributable to a single low-frequency variant (p.Asp563Gly; SKAT broad mask, conditional *p*=0.15). A second, previously-described, low-frequency variant *PAM* p.Ser539Trp^19^ is not represented on the exome array, and thus did not contribute to our analyses.

### Fine-mapping of coding variant association signals with T2D susceptibility

The present study has identified 40 distinct coding variant associations with T2D, but this information is not sufficient to determine that the variants themselves are causal for the disease. To assess the role of these coding variants in the context of regional genetic variation at the locus, we fine-mapped the distinct association signals using a European ancestry GWAS meta-analysis including 50,160 T2D cases and 465,272 controls (partially overlapping with the discovery samples), aggregated from 24 studies by the DIAGRAM Consortium. Each component GWAS had been imputed using suitable high density reference panels **(Methods, Supplementary Table 8**): (i) 22 GWAS were imputed up to the Haplotype Reference Consortium^20^; (ii) the UK Biobank GWAS was imputed to a merged reference panel from the 1000 Genomes Project (multi-ethnic, phase 3, October 2014 release)^21^ and the UK10K Project^9^; and (iii) the deCODE GWAS was imputed up to the deCODE Icelandic population-specific reference panel based on whole-genome sequence data^19^ (**Methods, Supplementary Table 8**). Distinct association signals were delineated before fine-mapping using approximate conditional analyses(**Methods, Supplementary Table 5**). We included 37 of the 40 identified coding variants in this fine-mapping analysis, excluding three that were not amenable to fine-mapping in the GWAS data sets we used: (i) the locus in the major histocompatibility complex because of the extended and complex structure of LD across the region, which complicates fine-mapping efforts; (ii) the East Asian specific *PAX4* p.Arg190His (rs2233580) signal, since the variant was not present in European ancestry GWAS; and (iii) *ZHX3* p.Asn310Ser (rs4077129) because the variant was only weakly associated with T2D in the GWAS data sets used for fine-mapping.

For each of the remaining 37 signals, we first constructed “functionally unweighted” credible variant sets which, at each locus, collectively account for 99% of the posterior probability (*π*_U_) of driving the association, based exclusively on the meta-analysis summary statistics^22^ **(Methods, Supplementary Table 9**). For each signal, we then calculated the total posterior probability attributed to coding variants (missense, in-frame indel, and splice region variants; **Figure 1, Supplementary Fig. 4 and 5**). Under this model, there were only two signals at which coding variants accounted for ≥80% of the posterior probability of association: *HNF4A* p.Thr139Ile (rs1800961, *π*_U_>0.999) and *RREB1* p. Asp1171Asn (rs9379084, *π*_U_=0.920). However, at other signals, including those for *GCKR* p.Pro446Leu and *SLC30A8* p.Arg276Trp, where robust empirical evidence has established their causal role^23,24^, the genetic evidence supporting coding variant causation was weak. This is because coding variants were typically in high LD (r^2^>0.9) with large numbers of non-coding variants, such that the posterior probabilities of association were distributed across many variants with broadly equivalent evidence for association.

**Figure 1.**
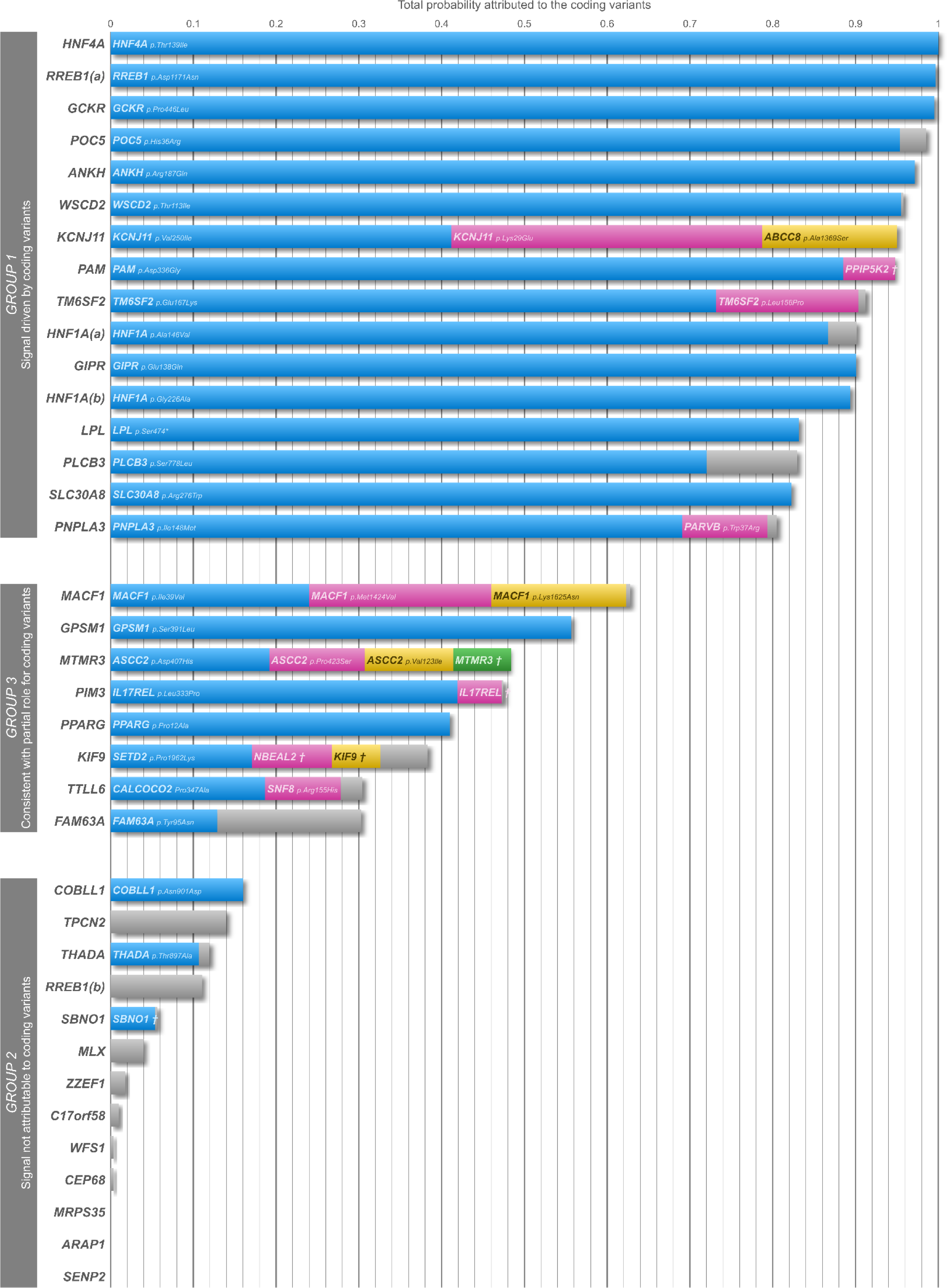
Posterior probabilities for coding variants across loci with annotation-informed priors. Fine-mapping of 37 distinct association signals was performed using European ancestry GWAS meta-analysis including 50,160 T2D cases and 465,272 controls. For each signal, we constructed a credible set of variants accounting for 99% of the posterior probability of driving the association, incorporating an “annotation informed” prior model of causality which “boosts” the posterior probability of driving the association signal that is attributed to coding variants. Each bar represents a signal with the total probability attributed to the coding variants within the 99% credible set plotted on the y-axis. When the probability (bar) is split across multiple coding variants (at least 0.05 probability attributed to a variant) at a particular locus, these are indicated by blue, pink, yellow, and green colours. The combined probability of the remaining coding variants are highlighted in grey. *RREB1*(a): *RREB1* p. Asp1171Asn; *RREB1*(b): *RREB1* p.Ser1499Tyr; *HNF1A*(a): *HNF1A* p.Ala146Val; *HNF1A*(b): *HNF1A* p.Ile75Leu; *PPIP5K2*†: *PPIP5K2* p.Ser1207Gly; *MTMR3*†: *MTMR3* p.Asn960Ser; *IL17REL*†: *IL17REL* p.Gly70Arg; *NBEAL2*†: *NBEAL2* p.Arg511Gly, *KIF9*†: *KIF9* p.Arg638Trp.

These functionally unweighted sets are based on genetic fine-mapping data alone, and do not account for the insight that coding variants are disproportionately represented amongst GWAS associations with complex traits^8,9^. To accommodate this knowledge, we extended these fine-mapping analyses by incorporating an “annotation informed” prior model of causality. We derived the priors from estimates of enrichment of association signals by sequence annotation from an analysis conducted by deCODE across 96 quantitative and 123 binary phenotypes^16^ (**Methods**). This model “boosts” the prior and hence the posterior probabilities (*π*_A_) coding variants. It also takes some account (in a tissue-non-specific way) of the GWAS enrichment of variants within enhancer elements (as assayed through DNase I hypersensitivity) as compared to non-coding variants mapping elsewhere. The annotation informed prior model generated smaller 99% credible sets across most signals, corresponding to fine-mapping at higher resolution (**Supplementary Table 9**).

As expected, the estimated contribution of coding variants was increased under the annotation informed model. At these 37 association signals, we could distinguish three broad patterns of causal relationships between coding variants and T2D risk.

### Group 1: T2D association signal is driven by coding variants

At 16 of the 37 distinct signals, coding variation accounted for >80% of the posterior probability of the association signal under the annotation informed model (**Figure 1, Table 2, Supplementary Table 9**). This posterior probability was attributed to a single coding variant at 12 signals and multiple coding variants at four. Reassuringly, group 1 signals confirmed coding variant causation for several loci at which functional studies (involving manipulation of the variant and/or effector gene) have reinforced genetic association data, establishing the role of *GCKR*, *PAM*, *SLC30A8*, and three variants in strong LD with each other at the *KCNJ11*-*ABCC8* locus (**Table 2**). T2D association signals at the 12 remaining signals (**Fig. 1, Supplementary Table 9**) had not previously been established to be driven by coding variation, but our fine-mapping analyses pointed to high probability causal coding variants after incorporating the annotation informed priors: these included *HNF4A*, *RREB1* (p. Asp1171Asn), *ANKH*, *WSCD2*, *POC5*, *TM6SF2*, *HNF1A* (p.Ala146Val and p.Ile75Leu), *GIPR*, *LPL*, *PLCB3*, and *PNPLA3* (**Table 2**). At several of these loci, independent evidence corroborates the causal role of the genes harbouring the associated coding variants with respect to T2D-risk. For example, rare coding mutations at *HNF1A* and *HNF4A* are causal for monogenic, early-onset forms of diabetes^25^; and at *TM6SF2* and *PNPLA3*, the coding variant concerned has been directly implicated in the development of non-alcoholic fatty liver disease (NAFLD)^26,27^.

**Table 2.**
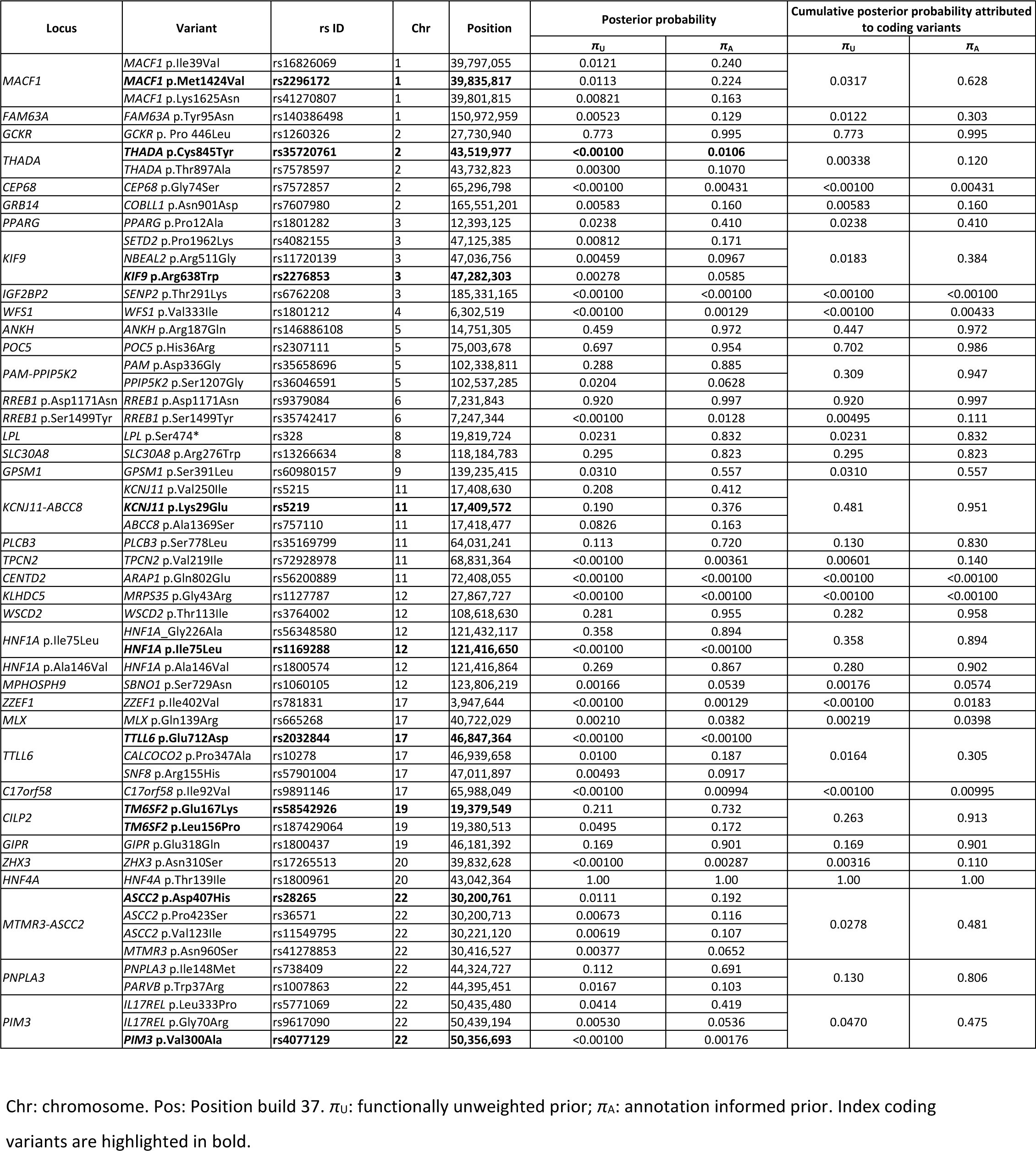
Posterior probabilities for coding variants within 99% credible set across loci with annotation informed and functionally unweighted prior based on fine-mapping analysis performed using 50,160 T2D cases and 465,272 controls of European ancestry.

The use of priors to capture the enrichment of coding variants seems a reasonable model, genome-wide. However, at any given locus, strong priors (especially for PTVs) might elevate to apparent causality, variants, which, on the basis of genetic fine-mapping alone, would have been excluded from a causal role. However, comparison of the annotation informed and functionally unweighted credible sets indicated that this was unlikely to be the case for any of the 16 association signals in group 1. For 11 of the 16 (*GCKR*, *PAM*, *KCNJ11*-*ABCC8*, *HNF4A*, *RREB1* [p. Asp1171Asn], *ANKH*, *POC5*, *TM6SF2*, *HNF1A* [p.Ala146Val], *PLCB3*, *PNPLA3*) the coding variant was the lead SNP in the fine-mapping analysis (**Table 2**), with the highest posterior probability of association, even under the functionally unweighted model. At *SLC30A8*, *WSCD2*, and *GIPR*, the coding variants had similar posterior probabilities of association as the lead non-coding SNPs under the functionally unweighted prior: *SLC30A8* p.Arg276Trp (rs13266634, *π*_U_0.295) and rs35859536, (*π*_U_=0.388); *WSCD2* p.Thr113Ile (rs3764002, *π*_U_=0.281) and rs1426371 (*π*_U_=0.475); and *GIPR* p.Glu138Gln (rs1800437, *π*_U_=0.169) and rs10423928 (*π*_U_=0.221). At these 14 signals therefore, the coding variants have either greater or equivalent posterior probabilities of association as the best flanking non-coding SNPs under the functionally unweighted model, but receive a boost in the posterior probability after accounting for variant annotation.

The situation is less clear at the *LPL* locus. Here, fine-mapping resolution is poor under the functionally unweighted prior, and the coding variant resides on an extended haplotype in strong LD with non-coding variants, some with higher posterior probabilities, such as rs74855321 (*π*_U_=0.0481) (compared to *LPL* p.Ser474* [rs328, *π*_U_=0.0231]). However, because *LPL* p.Ser474* is annotated as a PTV, it benefits from a substantially increased prior that is reflected in the annotation informed ranking. Ultimately, a decision on the causal role of any such variant must rest on the amalgamation of evidence from diverse sources including detailed functional evaluation of the coding variants, and of other variants with which they are in LD.

At the *HNF1A* p.Ile75Leu signal, the total posterior probability attributed to coding variants under the annotation informed prior was 0.894. However, the total probability was accounted for by p.Gly226Ala (rs56348580,*π*_A_=0.894), a variant missing from the exome array. Conversely, the posterior probability attributed to the index coding variant p.Ile75Leu (rs1169288) was <0.001, although this may reflect its absence from most commercial GWAS arrays and low-quality imputation. Fine-mapping analyses conducted using the Metabochip^10^, on which *HNF1A* p.Ile75Leu, as well as many local non-coding variants, are directly typed, demonstrates that these two coding variants are likely to be driving distinct association signals at this locus, and are consistent with our observations from the exome array. Direct genotyping or sequencing will be required to fully disentangle the relationships between the various coding variants at this signal. However, the established role of rare coding variants in *HNF1A* with respect to monogenic forms of diabetes leaves little doubt concerning the role of this gene as the effector transcript at this locus.

### Group 2: T2D association signals are not attributable to coding variants

At 13 of the 37 distinct signals, coding variation accounted for <20% of the posterior probability of driving the association, even after applying the annotation informed prior model that boosts coding variant posterior probabilities. These signals are likely to be driven by local non-coding variation and mediated through regulation of gene expression. Five of these signals (*TPCN2*, *MLX*, *ZZEF1*, C17orf58, and *CEP68*) represent novel T2D-association signals identified in the exome-focused analysis. On the basis of the exome-array discoveries, it would have been natural to consider the named genes at these, and the other loci in this group, as strong candidates for mediation of their respective association signals. However, the fine-mapping analyses indicate that these coding variants cannot provide useful mechanistic inference given their low posterior probability of driving association (**Fig. 1, Table 2**).

The coding variant association at the *CENTD2* (*ARAP1*) locus is a case-in-point. The association with the p.Gln802Glu variant in *ARAP1* (rs56200889, ***p***_TE_=4.8×10^−8^ but *π*_A_<0.001 in the annotation informed analysis) is clearly seen in the fine-mapping analysis to be secondary to a substantially stronger non-coding association signal involving a cluster of variants including rs11603334 (***p***_TE_=9.5×10^−18^, *π*_A_=0.0692) and rs1552224 (***p***_TE_=2.5×10^−17^, *π*_A_=0.0941). The identity of the effector transcript at this locus has been the subject of considerable investigation, and some early studies used islet expression data to highlight *ARAP1* as the strongest candidate^28^. However, a more recent study integrating studies of human islet genomics and murine gene knockouts points firmly towards *STARD10* as the gene mediating the GWAS signal, consistent with the reassignment of the *ARAP1* coding variant association as irrelevant to biological inference at this locus^29^.

Whilst, at these loci, the coding variant associations appear to be “false leads”, this does not necessarily exclude the genes concerned from a causal role. At *WFS1* for example, coding variants too rare to be visible to the array-based analyses we performed, and statistically independent of the common p.Val333Ile variant we detected, cause an early-onset form of diabetes that makes *WFS1* the strongest local candidate for T2D predisposition. An additional example emerged from analyses to uncover causal non-coding mechanisms at these loci. When we extracted variants with posterior probability >0.05 from the fine-mapping analysis and searched for *cis*-eQTL signals across multiple tissues (including islets) using public resources (**Supplementary Table 10**)^30,31^, the only significant signals implicated *CEP68*. Non-coding variants in the 99% credible set were associated with *CEP68* expression across multiple tissues including visceral adipose tissue. This highlights *CEP68* as a candidate effector transcript at this locus, even though the coding variant signal at p.Gly74Ser is clearly not causal.

### Group 3: Fine-mapping data consistent with partial role for coding variants

At eight of the 37 distinct signals, the total posterior probability attributable to coding variation in the annotation informed analyses was between 20% and 80%. At these signals, the evidence is consistent with “partial” contributions from coding variants, although the precise inference is likely to be locus-specific, dependent on subtle variations in LD, imputation accuracy, and the extent to which the global priors accurately represent the functional impact of the specific variants concerned.

This group includes *PPARG* for which independent evidence corroborates the causal role of this specific effector transcript with respect to T2D risk. *PPARG* encodes the target of antidiabetic thiazolidinedione drugs and is known to harbour rare coding variants that are causal for lipodystrophy and insulin resistance, both conditions highly relevant to T2D. The common variant association signal at this locus has generally been attributed to the p.Pro12Ala coding variant (rs1801282) although confirmation that this variant has an empirical impact on PPARG function has been difficult to obtain^32–34^. In the functionally unweighted analysis, p.Pro12Ala had an unimpressive posterior probability of being causal (*π*_U_=0.0238); after including annotation informed priors, the same variant emerged with the highest posterior probability (*π*_A_=0.410), although the 99% credible set included 19 noncoding variants, spanning 67kb (**Supplementary Table 9**). These credible set variants included rs4684847 (*π*_A_=0.00891), at which the T2D-associated allele has been reported to impact *PPARG2* expression and insulin sensitivity by altering binding of the homeobox transcription factor PRRX1^35^. These data are consistent with a model whereby regulatory variants contribute to the mechanisms through which the T2D GWAS signal impacts PPARG activity (in combination with, or potentially to the exclusion of, p.Pro12Ala). Future improvements in functional annotation for regulatory variants (gathered from relevant tissues and cell types) should provide increasingly granular priors that can be used to fine-tune assignment of causality at loci such as this.

### Functional impact of coding alleles

In other contexts, the functional impact of coding alleles is correlated with: (i) variant-specific features, including measures of conservation and predicted impact on protein structure; and (ii) gene-specific features such as extreme selective constraints as quantified by the intolerance to functional variation^36^. To determine whether similar measures could capture information pertinent to T2D causation, we compared coding variants falling into the different fine-mapping groups for a variety of measures including MAF, Combined Annotation Dependent Depletion score (CADD-score)^37^, and loss-of-function (LoF)-intolerance metric, pLI^36^ (**Methods, Fig. 2**). As noted previously^37^, CADD-score and MAF exhibit a negative correlation (Pearson′s correlation r=-0.44, *p*=0.0033) across association signals. Variants from group 1 had significantly higher CADD-scores than those in group 2 (*p*=0.0031, by Kolmogorov-Smirnov test). With the exception of the variants at *KCNJ11*-*ABCC8* and *GCKR*, all group 1 coding variants considered likely to be driving T2D association signals have CADD-score ≥20 (i.e. the 1% most deleterious variants predicted across the human genome). On this basis, we would predict that the East-Asian specific coding variant at *PAX4*, for which the fine-mapping data were not informative, is also likely causal for T2D.

**Figure 2.**
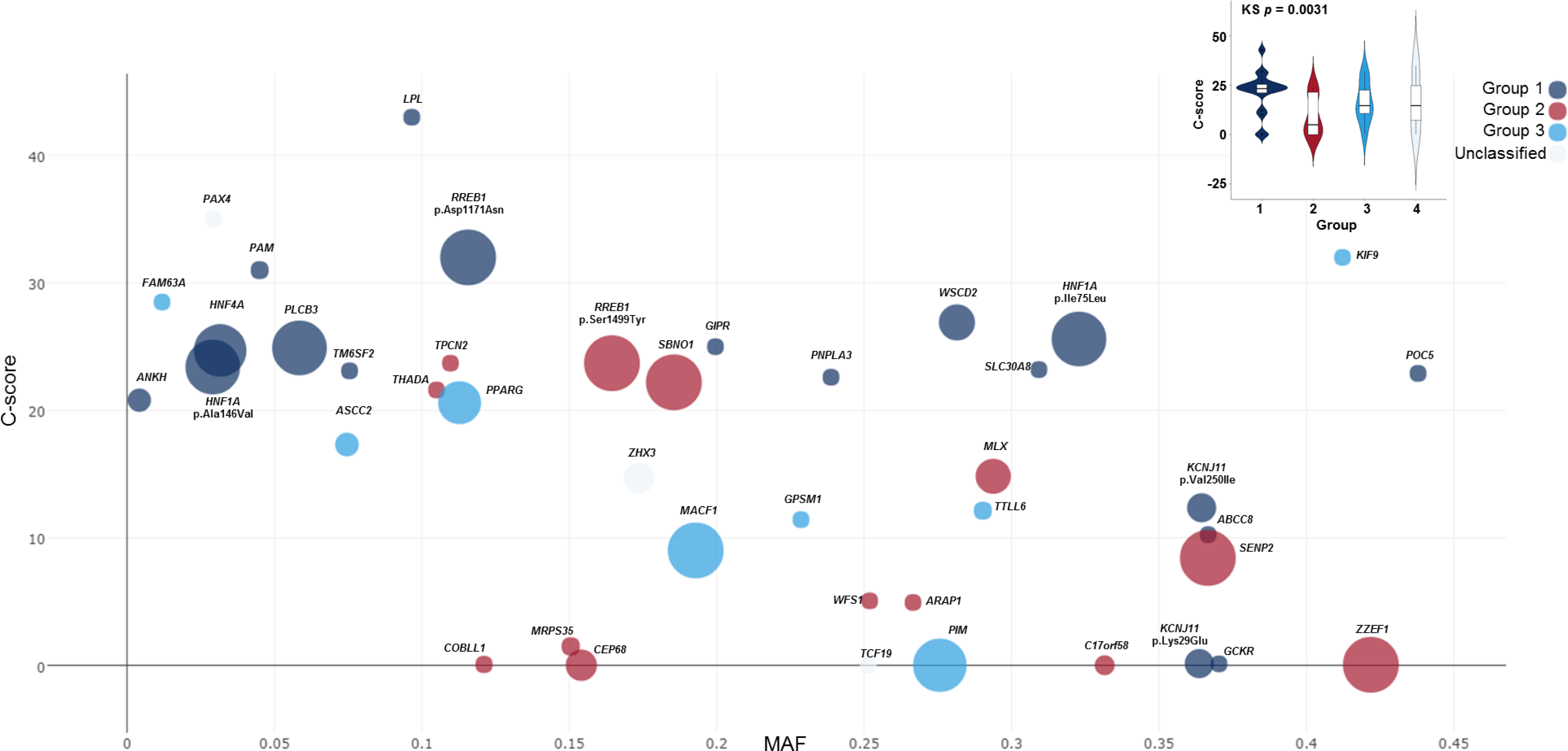
Plot of measures of variant-specific and gene-specific features of distinct coding signals to access the functional impact of coding alleles. Each point represents a coding variant with the minor allele frequency (MAF) plotted on the x-axis and the Combined Annotation Dependent Depletion score (CADD-score) plotted on the y-axis. Size of each point varies with the measure of intolerance of the gene to loss of function variants (pLI) and the colour represents the fine-mapping group each variant is assigned to. Group 1: signal is driven by coding variant. Group 2: signal attributable to non-coding variants. Group 3: consistent with partial role for coding variants. Unclassified category includes *PAX4* and signal at *TCF19* within the MHC region where we did not perform fine-mapping. The distribution of CADD-score between different groups is shown in the inset.

### Novel non-coding association signals for T2D susceptibility

Whilst the exome array primarily encompasses variation that alters protein function, it also incorporates previously reported non-coding lead SNPs from GWAS for a range of complex human phenotypes, including metabolic traits that may also impact T2D susceptibility. We detected novel significant (*p*<5×10^−8^) associations with T2D status for 20 non-coding variants at 15 loci. Three of these (*POC5*, *LPL*, and *BPTF*) overlap with novel coding signals reported here (**Supplementary Table 11**). Lead SNPs at these loci had been included on the exome array as GWAS tags for association signals with metabolic traits including central and overall obesity, lipid levels, coronary heart disease, venous thromboembolism, and menarche, but here were demonstrated to be genome-wide significant for T2D as well (**Supplementary Table 11**). If instead of the conventional genome-wide significance threshold, we adopted a more conservative reweighted Bonferroni threshold for non-coding variants of *p*<9.5×10^−9^, balancing the less stringent threshold used for defining significance for coding variants, 10 of the 15 loci remained associated with T2D.

### T2D loci and physiological classification

The development of T2D involves dysfunction of multiple mechanisms. Systematic analyses of the physiological effects of known T2D risk variants have improved understanding of the key intermediate processes involved in the disease and the mechanisms through which many of those variants exert their primary impact on disease risk^38^. We obtained association summary statistics for a range of metabolic traits and other outcomes for 94 T2D-associated index variants (again restricted to those represented on the exome array) representing the 40 distinct coding signals (not necessarily causal) and 54 distinct non-coding signals (including 12 novel and 42 previously reported non-coding GWAS lead SNPs). We applied hierarchical clustering techniques (**Methods**) to generate multi-trait association patterns, and were able to allocate 71 of the 94 loci to one of three categories (**Supplementary Fig. 6, Supplementary Table 12**). The first category, comprising of nine T2D-risk loci with strong BMI and dyslipidemia phenotypic associations, included three of the novel coding signals: *PNPLA3*, which showed strong association with dyslipidemia, and *POC5* and *BPTF* where the risk allele was associated with increased BMI (**Supplementary Fig. 6, Supplementary Table 12**). The T2D associations at both *POC5* and *BPTF* were substantially attenuated (at least by 2-fold decrease in −log10p) after adjusting for BMI (**Table 1, Supplementary Table 3, Supplementary Fig. 7**), indicating that the impact on T2D risk is likely mediated by a primary effect through increased adiposity. *PNPLA3* and *POC5* are established NAFLD^26^ and BMI^6^ loci, respectively. The second category included 39 loci at which the multi-trait profiles indicated a primary effect on insulin secretion. This category included four of the novel coding variant signals (*ANKH*, *ZZEF1*, *TTLL6*, and *ZHX3*). The third category encompassed 23 loci with primary effects on insulin action including coding variant associations at the *KIF9*, *PLCB3*, *CEP68*, *TPCN2*, *FAM63A*, and *PIM3* loci. For most variants in this category, the T2D-risk allele was associated with lower BMI, and T2D association signals were more pronounced after adjustment for BMI. At a subset of these loci, including *KIF9* and *PLCB3*, T2D-risk alleles were associated with markedly higher waist-hip ratio and lower body fat percentage, indicating that their mechanism of action likely reflects limitations in storage capacity of peripheral adipose tissue^39^.

## DISCUSSION

The present study adds to mounting evidence constraining the contribution of lower-frequency variants to T2D risk. Although the exome array interrogates only a subset of the universe of coding variants, it does capture the majority of low-frequency (MAF>0.5%) coding variants in European populations. The substantial increase in sample size in the present study over our previous effort^12^ (effective sample sizes of 228,825 and 82,758, respectively), provides more robust enumeration of the effect size distribution in this low-frequency variant range, and indicates that previous analyses are likely to have, if anything, overestimated the contribution of low-frequency variants to T2D-risk.

The present study is less informative regarding rare variants. These are sparsely captured on the exome array. In addition, the combination of greater regional diversity in rare allele distribution and the enormous sample sizes required to detect many rare variant associations (which would require meta-analysis of data from multiple diverse populations) acts against their detection. We note that our complementary genome and exome sequence analyses have thus far failed to register strong evidence for a substantial rare variant component to T2D-risk^12^. It is therefore highly unlikely that rare variants missed in our analyses are causal for any of the common or low-frequency variant associations we have detected and fine-mapped in this analysis. On the other hand, it <u>is</u> probable that rare coding alleles, with associations that are statistically independent of the common variant signals we have examined and only revealed through sequence based analyses, will provide additional clues to the most likely effector transcripts at some of these loci (*WFS1* represents such a locus).

Once a coding variant association is detected, it is natural to assume a causal connection between that variant, the gene in which it sits, and the disease or phenotype of interest. Whilst such assignments may be robust for many rare protein-truncating alleles, we demonstrate that this implicit assumption is often inaccurate, particularly for associations attributable to common, missense variants. A third of the coding variant associations we detected were, when assessed in terms of regional LD, highly unlikely to be causal. At these loci, the genes within which they reside are consequently deprived of their implied connection to disease risk, and attention redirected towards nearby non-coding variants and their impact on regional gene expression. As a group, coding variants we assign as causal are predicted to have a more deleterious impact on gene function than those that we exonerate, but, as in other settings, coding annotation methods lack both sensitivity and specificity. Besides, it is worth emphasising that empirical evidence that the associated coding allele is “functional” (i.e. that it can be shown to influence function of its cognate gene in some experimental assay) provides limited reassurance that the coding variant is responsible for the T2D association, unless that specific perturbation of gene function can itself be plausibly linked to the disease phenotype.

Our fine-mapping analyses make use of the observation that coding variants are globally enriched across GWAS signals^8,9,16^ with greater prior probability of causality assigned to those with more severe impact on biological function. We assigned diminished priors to non-coding variants, with lowest support for those mapping outside of DHS. The extent to which our findings corroborate previous assignments of causality (often backed up by detailed, disease appropriate functional assessment and orthogonal causal evidence) suggests that even these sparse annotations provide valuable information to guide target validation efforts. Nevertheless, we recognise that there are inevitable limits to the extrapolation of these broad-brush genome-wide enrichments to individual loci, and expect that improvements in functional annotation for both coding and regulatory variants, particularly when these are gathered from trait-relevant tissues and cell types, will provide more granular, trait-specific priors to fine-tune assignment of causality within associated regions. These will motivate target validation efforts that benefit from synthesis of both coding and regulatory routes to gene perturbation. It also needs to be acknowledged that, in the absence of whole genome sequencing data on sample sizes comparable to those we have examined here, imperfections arising from the imputation process have the potential to confound fine-mapping precision at some loci, and that robust inference of causation will inevitably depend on integration of diverse sources of genetic, genomic and functional data.

The term “smoking gun” has often been used to describe the potential of functional coding variants to provide causal inference with respect to pathogenetic mechanisms^40^: our study provides a timely reminder that, even when a suspect with a smoking gun is found at the scene of a crime, it should not automatically be assumed that they fired the fatal bullet.

## DATA AVAILABILITY STATEMENT

Summary level data from the exome array component of this project will be made available at the DIAGRAM consortium website http://diagram-consortium.org/ and Accelerating Medicines Partnership T2D portal http://www.type2diabetesgenetics.org/.

## ACKNOWLEDGMENTS

A full list of acknowledgments appears in the Supplementary Information. Part of this work was conducted using the UK Biobank resource.

## AUTHOR CONTRIBUTIONS

### Project co-ordination

A.Mahajan, A.P.M., J.I.R., M.I.M.

### Core analyses and writing

A.Mahajan, J.W., S.M.W, W.Zhao, N.R.R., A.Y.C., W.G., H.K., R.A.S., I.Barroso, T.M.F., M.O.G., J.B.M., M.Boehnke, D.S., A.P.M., J.I.R., M.I.M.

### Statistical Analysis in individual studies

A.Mahajan, J.W., S.M.W., W.Zhao, N.R.R., A.Y.C., W.G., H.K., D.T., N.W.R., X.G., Y.Lu, M.Li, R.A.J., Y.Hu, S.Huo, K.K.L., W.Zhang, J.P.C., B.P., J.Flannick, N.G., V.V.T., J.Kravic, Y.J.K., D.V.R., H.Y., M.M.-N., K.M., R.L.-G., T.V.V., J.Marten, J.Li, A.V.S., P.An, S.L., S.G., G.M., A.Demirkan, J.F.T., V.Steinthorsdottir, M.W., C.Lecoeur, M.Preuss, L.F.B., P.Almgren, J.B.-J., J.A.B., M.Canouil, K.-U.E., H.G.d.H., Y.Hai, S.Han, S.J., F.Kronenberg, K.L., L.A.L., J.-J.L., H.L., C.-T.L., J.Liu, R.M., K.R., S.S., P.S., T.M.T., G.T., A.Tin, A.R.W., P.Y., J.Y., L.Y., R.Y., J.C.C., D.I.C., C.v.D., J.Dupuis, P.W.F., A.Köttgen, D.M.-K., N.Soranzo, R.A.S., A.P.M.

### Genotyping

A.Mahajan, N.R.R., A.Y.C., Y.Lu, Y.Hu, S.Huo, B.P., N.G., R.L.-G., P.An, G.M., E.A., N.A., C.B., N.P.B., Y.-D.I.C., Y.S.C., M.L.G., H.G.d.H., S.Hackinger, S.J., B.-J.K., P.K., J.Kriebel, F.Kronenberg, H.L., S.S.R., K.D.T., E.B., E.P.B., P.D., J.C.F., S.R.H., C.Langenberg, M.A.P., F.R., A.G.U., J.C.C., D.I.C., P.W.F., B.-G.H., C.H., E.I., S.L.K., J.S.K., Y.Liu, R.J.F.L., N.Soranzo, N.J.W., R.A.S., T.M.F., A.P.M., J.I.R., M.I.M.

### Cross-trait lookups in unplublished data

S.M.W., A.Y.C., Y.Lu, M.Li, M.G., H.M.H., A.E.J.,D. J.L., E.M., G.M.P., H.R.W., S.K., C.J.W.

### Phenotyping

Y.Lu, Y.Hu, S.Huo, P.An, S.L., A.Demirkan, S.Afaq, S.Afzal, L.B.B., A.G.B., I.Brandslund, C.C., S.V.E., G.G., V.Giedraitis, A.T.-H., M.-F.H., B.I., M.E.J., T.J., A.Käräjämäki, S.S.K., H.A.K., P.K., F.Kronenberg, B.L., H.L., K.-H.L., A.L., J.Liu, M.Loh, V.M., R.M.-C., G.N., M.N., S.F.N., I.N., P.A.P., W.R., L.R., O.R., S.S., E.S., K.S.S., A.S., B.T., A.Tönjes, A.V., D.R.W.,H.B., E.P.B., A.Dehghan, J.C.F., S.R.H., C.Langenberg, A.D.Morris, R.d.M., M.A.P., A.R., P.M.R., F.R.R., V.Salomaa, W.H.-H.S., R.V., J.C.C., J.Dupuis, O.H.F., H.G., B.-G.H., T.H., A.T.H., C.H.,S.L.K., J.S.K., A.Köttgen, L.L., Y.Liu, R.J.F.L., C.N.A.P., J.S.P., O.P., B.M.P., M.B.S., N.J.W.,T.M.F., M.O.G.

### Individual study design and principal investigators

N.G., P.An, B.-J.K., P.Amouyel, H.B., E.B.,E. P.B., R.C., F.S.C., G.D., A.Dehghan, P.D., M.M.F., J.Ferrières, J.C.F., P.Frossard, V.Gudnason, T.B.H., S.R.H., J.M.M.H., M.I., F.Kee, J.Kuusisto, C.Langenberg, L.J.L., C.M.L., S.M., T.M., O.M., K.L.M., M.M., A.D.Morris, A.D.Murray, R.d.M., M.O.-M., K.R.O., M.Perola, A.P., M.A.P., P.M.R., F.R., F.R.R., A.H.R., V.Salomaa, W.H.-H.S., R.S., B.H.S., K.Strauch, A.G.U., R.V., M.Blüher, A.S.B., J.C.C., D.I.C., J.Danesh, C.v.D., O.H.F., P.W.F., P.Froguel, H.G., L.G., T.H., A.T.H., C.H., E.I., S.L.K., F.Karpe, J.S.K., A.Köttgen, K.K., M.Laakso, X.L., L.L., Y.Liu, R.J.F.L., J.Marchini, A.Metspalu, D.M.-K., B.G.N., C.N.A.P., J.S.P., O.P., B.M.P., R.R., N.Sattar, M.B.S., N.Soranzo, T.D.S., K.Stefansson, M.S., U.T., T.T., J.T., N.J.W., J.G.W., E.Z., I.Barroso, T.M.F., J.B.M., M.Boehnke, D.S., A.P.M., J.I.R., M.I.M.

## MATERIALS & CORRESPONDENCE

Correspondence and requests for materials should be addressed to.

## ONLINE METHODS

### Ethics statement

All human research was approved by the relevant institutional review boards, and conducted according to the Declaration of Helsinki. All participants provided written informed consent.

### Derivation of significance thresholds

We considered five categories of annotation^16^ of variants on the exome array in order of decreasing effect on biological function: (1) PTVs (stop-gain and stop-loss, frameshift indel, donor and acceptor splice-site, and initiator codon variants, *n*_1_=8,388); (2) moderate-impact variants (missense, in-frame indel, and splice region variants, *n*_2_=216,114); (3) low-impact variants (synonymous, 3′ and 5′ UTR, and upstream and downstream variants, *n*_3_=8,829); (4) other variants mapping to DNase I hypersensitive sites in any of 217 cell types^8^ (DHS, *n*_4_=3,561); and (5) other variants not mapping to DHS (*n*_5_=10,578). To account for the greater prior probability of causality for variants with greater effect on biological function, we determined a weighted Bonferroni-corrected significance threshold on the basis of reported enrichment^16^, denoted *w_i_*, in each annotation category, *i*: *W*_1_=165; *W*_2_=33; *W*_3_=3; *W*_4_=1.5; *W*_5_=0.5. For coding variants (annotation categories 1 and 2):

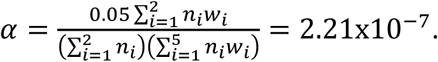

We note that this threshold is similar to a simple Bonferroni correction for the total number of coding variants on the array, which would yield:

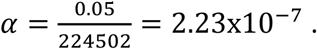

For non-coding variants (annotation categories 3, 4 and 5) the weighted Bonferroni-corrected significance threshold is:

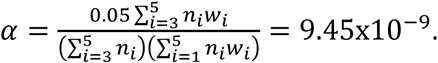

### DISCOVERY: Exome-array study-level analyses

Within each study, genotype calling and quality control were undertaken according to protocols developed by the UK Exome Chip Consortium or the CHARGE central calling effort^41^ (**Supplementary Table 1**). Within each study, variants were then excluded for the following reasons: (i) not mapping to autosomes or X chromosome; (ii) multi-allelic and/or insertion-deletion; (iii) monomorphic; (iv) call rate <99%; or (v) exact *p*<10^−4^ for deviation from Hardy-Weinberg equilibrium (autosomes only).

We tested association of T2D with each variant in a linear mixed model, implemented in RareMetalWorker^17^, using a genetic relationship matrix (GRM) to account for population structure and relatedness. For participants from family-based studies, known relationships were incorporated directly in the GRM. For founders and participants from population-based studies, the GRM was constructed from pair-wise identity by descent (IBD) estimates based on LD pruned (*r*^2^<0.05) autosomal variants with MAF≥1% (across cases and controls combined), after exclusion of those in high LD and complex regions^42,43^, and those mapping to established T2D loci. We considered additive, dominant, and recessive models for the effect of the minor allele, adjusted for age and sex (where appropriate) and additional study-specific covariates (**Supplementary Table 2**). Analyses were also performed with and without adjustment for BMI (where available Supplementary Table 2).

For single-variant association analyses, variants with minor allele count ≤10 in cases and controls combined were excluded. Association summary statistics for each analysis were corrected for residual inflation by means of genomic control^44^, calculated after excluding variants mapping to established T2D susceptibility loci. For gene-based analyses, we made no variant exclusions on the basis of minor allele count.

### DISCOVERY: Exome-sequence analyses

We used summary statistics of T2D association from analyses conducted on 8,321 T2D cases and 8,421 controls across different ancestries, all genotyped using exome sequencing. Details of samples included, sequencing, and quality control are described elsewhere^12,15^ (http://www.type2diabetesgenetics.org/). Samples were subdivided into 15 sub-groups according to ancestry and study of origin. Each subgroup was analysed independently, with sub-group specific principal components and genetic relatedness matrices. Association tests were performed with both a linear mixed model, as implemented in EMMAX^45^, using covariates for sequencing batch, and the Firth test, using covariates for principal components and sequencing batch. Related samples were excluded from the Firth analysis but maintained in the linear mixed model analysis. Variants were then filtered from each sub-group analysis, according to call rate, differential case-control missing-ness, or deviation from Hardy-Weinberg equilibrium (as computed separately for each sub-group). Association statistics were then combined via a fixed-effects inverse-variance weighted meta-analysis, at both the level of ancestry as well as across all samples. P-values were taken from the linear mixed model analysis, while effect sizes estimates were taken from the Firth analysis. Analyses were performed with and without adjustment for BMI. From exome sequence summary statistics, we extracted variants passing quality control and present on the exome array.

### DISCOVERY: GWAS analyses

The UK Biobank is a large detailed prospective study of more than 500,000 participants aged 40-69 years when recruited in 2006-2010^13^. Prevalent T2D status was defined using self-reported medical history and medication in UK Biobank participants^46^. Participants were genotyped with the UK Biobank Axiom Array or UK BiLEVE Axiom Array, and quality control and population structure analyses were performed centrally at UK Biobank. We defined a subset of “white European” ancestry samples (n=120,286) as those who both self-identified as white British and were confirmed as ancestrally “Caucasian” from the first two axes of genetic variation from principal components analysis. Imputation was also performed centrally at UK Biobank for the autosomes only, up to a merged reference panel from the 1000 Genomes Project (multiethnic, phase 3, October 2014 release)^21^ and the UK10K Project^9^. We used SNPTESTv2.5^47^ to test for association of T2D with each SNP in a logistic regression framework under an additive model, and after adjustment for age, sex, six axes of genetic variation, and genotyping array as covariates. Analyses were performed with and without adjustment for BMI, after removing related individuals.

GERA is a large multi-ethnic population-based cohort, created for investigating the genetic and environmental basis of age-related diseases [dbGaP phs000674.p1]. T2D status is based on ICD-9 codes in linked electronic medical health records, with all other participants defined as controls. Participants have previously been genotyped using one of four custom arrays, which have been designed to maximise coverage of common and low-frequency variants in non-Hispanic white, East Asian, African American, and Latino ethnicities^48,49^. Methods for quality control have been described previously^14^. Each of the four genotyping arrays were imputed separately, up to the 1000 Genomes Project reference panel (autosomes, phase 3, October 2014 release; X chromosome, phase 1, March 2012 release) using IMPUTEv2.3^50,51^. We used SNPTESTv2.5^47^ to test for association of T2D with each SNP in a logistic regression framework under an additive model, and after adjustment for sex and nine axes of genetic variation from principal components analysis as covariates. BMI was not available for adjustment in GERA.

For UK Biobank and GERA, we extracted variants passing standard imputation quality control thresholds (IMPUTE info≥0.4)^52^ and present on the exome array. Association summary statistics under an additive model were corrected for residual inflation by means of genomic control^44^, calculated after excluding variants mapping to established T2D susceptibility loci: GERA (λ=1.097 for BMI unadjusted analysis) and UK Biobank (λ=1.043 for BMI unadjusted analysis, λ=1.056 for BMI adjusted analysis).

### DISCOVERY: Single-variant meta-analysis

We aggregated association summary statistics under an additive model across studies, with and without adjustment for BMI, using METAL^53^: (i) effective sample size weighting of *Z*-scores to obtain *p*-values; and (ii) inverse variance weighting of log-odds ratios. For exome-array studies, allelic effect sizes and standard errors obtained from the RareMetalWorker linear mixed model were converted to the log-odds scale prior to meta-analysis to correct for case-control imbalance^54^.

The European-specific meta-analyses aggregated association summary statistics from a total of 48,286 cases and 250,671 controls from: (i) 33 exome-array studies of European ancestry; (ii) exome-array sequence from individuals of European ancestry; and (iii) GWAS from UK Biobank. Note that non-coding variants represented on the exome array were not available in exome sequence. The European-specific meta-analyses were corrected for residual inflation by means of genomic control^44^, calculated after excluding variants mapping to established T2D susceptibility loci: λ=1.091 for BMI unadjusted analysis and λ=1.080 for BMI adjusted analysis.

The trans-ethnic meta-analyses aggregated association summary statistics from a total of 81,412 cases and 370,832 controls across all studies (51 exome array studies, exome sequence, and GWAS from UK Biobank and GERA), irrespective of ancestry. Note that noncoding variants represented on the exome array were not available in exome sequence. The trans-ethnic meta-analyses were corrected for residual inflation by means of genomic control^44^, calculated after excluding variants mapping to established T2D susceptibility loci: λ=1.073 for BMI unadjusted analysis and λ=1.068 for BMI adjusted analysis. Heterogeneity in allelic effect sizes between exome-array studies contributing to the trans-ethnic metaanalysis was assessed by Cochran′s *Q* statistic^55^.

### DISCOVERY: Detection of distinct association signals

Conditional analyses were undertaken to detect association signals by inclusion of index variants and/or tags for previously reported non-coding GWAS lead SNPs as covariates in the regression model at the study level. Within each exome-array study, approximate conditional analyses were undertaken under a linear mixed model using RareMetal^17^, which uses score statistics and the variance-covariance matrix from the RareMetalWorker single-variant analysis to estimate the correlation in effect size estimates between variants due to LD. Study-level allelic effect sizes and standard errors obtained from the approximate conditional analyses were converted to the log-odds scale to correct for case-control imbalance^54^. Within each GWAS, exact conditional analyses were performed under a logistic regression model using SNPTESTv2.5^47^. GWAS variants passing standard imputation quality control thresholds (IMPUTE info≥0.4)^52^ and present on the exome array were extracted for meta-analysis.

Association summary statistics were aggregated across studies, with and without adjustment for BMI, using METAL^53^: (i) effective sample size weighting of *Z*-scores to obtain *p*-values; and (ii) inverse variance weighting of log-odds ratios.

We defined novel loci as mapping >500kb from a previously reported lead GWAS SNP. We performed conditional analyses where a novel signal mapped close to a known GWAS locus, and the lead GWAS SNP at that locus is present (or tagged) on the exome array (**Supplementary Table 5**).

### DISCOVERY: Non-additive association models

For exome-array studies only, we aggregated association summary statistics under recessive and dominant models across studies, with and without adjustment for BMI, using METAL^53^: (i) effective sample size weighting of *Z*-scores to obtain *p*-values; and (ii) inverse variance weighting of log-odds ratios. Allelic effect sizes and standard errors obtained from the RareMetalWorker linear mixed model were converted to the log-odds scale prior to meta-analysis to correct for case-control imbalance^54^. The European-specific meta-analyses aggregated association summary statistics from a total of 41,066 cases and 136,024 controls from 33 exome-array studies of European ancestry. The European-specific meta-analyses were corrected for residual inflation by means of genomic control^44^, calculated after excluding variants mapping to established T2D susceptibility loci: λ=1.076 and λ=1.083 for BMI unadjusted analysis, under recessive and dominant models respectively, and λ=1.081 and λ=1.062 for BMI adjusted analysis, under recessive and dominant models respectively. The trans-ethnic meta-analyses aggregated association summary statistics from a total of 58,425 cases and 188,032 controls across all exome-array studies, irrespective of ancestry. The trans-ethnic meta-analyses were corrected for residual inflation by means of genomic control^44^, calculated after excluding variants mapping to established T2D susceptibility loci: λ=1.041 and λ=1.071 for BMI unadjusted analysis, under recessive and dominant models respectively, and λ=1.031 and λ=1.063 for BMI adjusted analysis, under recessive and dominant models respectively.

### DISCOVERY: Gene-based meta-analyses

For exome-array studies only, we aggregated association summary statistics under an additive model across studies, with and without adjustment for BMI, using RareMetal^17^. This approach uses score statistics and the variance-covariance matrix from the RareMetalWorker single-variant analysis to estimate the correlation in effect size estimates between variants due to LD. We performed gene-based analyses using a burden test (assuming all variants have same direction of effect on T2D susceptibility) and SKAT (allowing variants to have different directions of effect on T2D susceptibility). We used two previously defined filters for annotation and MAF^18^ to define group files: (i) strict filter, including 44,666 variants; and (ii) broad filter, including all variants from the strict filter, and 97,187 additional variants.

We assessed the contribution of each variant to gene-based signals by performing approximate conditional analyses. We repeated RareMetal analyses for the gene, excluding each variant in turn from the group file, and compared the strength of the association signal.

### Fine-mapping of coding variant association signals with T2D susceptibility

We defined a locus as mapping 500kb up- and down-stream of each index coding variant (**Supplementary Table 5**), excluding the MHC. Our fine-mapping analyses aggregated association summary statistics from 24 GWAS incorporating 50,160 T2D cases and 465,272 controls of European ancestry from the DIAGRAM Consortium (**Supplementary Table 8**). Each GWAS was imputed using miniMAC^12^ or IMPUTEv2^50,51^ up to reference panels from the Haplotype Reference Consortium^20^, the 1000 Genomes Project (multi-ethnic, phase 3, October 2014 release)^21^ and the UK10K Project^9^, or population-specific whole-genome sequence data^19^ (**Supplementary Table 8**). Association with T2D susceptibility was tested for each remaining variant using logistic regression, adjusting for age, sex, and study-specific covariates, under an additive genetic model. Analyses were performed with and without adjustment for BMI. For each study, variants with minor allele count<5 (in cases and controls combined) or those with imputation quality r2-hat<0.3 (miniMAC) or proper-info<0.4 (IMPUTE2) were removed. Association summary statistics for each analysis were corrected for residual inflation by means of genomic control^44^, calculated after excluding variants mapping to established T2D susceptibility loci

We aggregated association summary statistics across studies, with and without adjustment for BMI, in a fixed-effects inverse variance weighted meta-analysis, using METAL^53^. The BMI unadjusted meta-analysis was corrected for residual inflation by means of genomic control (λ=1.012)^44^, calculated after excluding variants mapping to established T2D susceptibility loci. No adjustment was required for BMI adjusted meta-analysis (λ=0.994). From the meta-analysis, variants were extracted that were present on the HRC panel and reported in at least 50% of total effective sample size.

To delineate distinct association signals in four regions, we undertook approximate conditional analyses, implemented in GCTA^56^, to adjust for the index coding variants and non-coding lead GWAS SNPs: (i) *RREB1* p. Asp1171Asn (rs9379084), p.Ser1499Tyr (rs35742417), and rs9505118; (ii) *HNF1A* p.Ile75Leu (rs1169288) and p.Ala146Val (rs1800574); (iii) *GIPR* p.Glu318Gln (rs1800437) and rs8108269; and (iv) *HNF4A* p.Thr139Ile (rs1800961) and rs4812831. We made use of summary statistics from the fixed-effects meta-analyses (BMI unadjusted for *RREB1*, *HNF1A*, and *HNF4A*, and BMI adjusted for *GIPR* as this signal was only seen in BMI adjusted analysis) and genotype data from 5,000 random individuals of European ancestry from the UK Biobank, as reference for LD between genetic variants across the region.

For each association signal, we first calculated an approximate Bayes′ factor^57^ in favour of association on the basis of allelic effect sizes and standard errors from the metaanalysis. Specifically, for the *j*th variant,

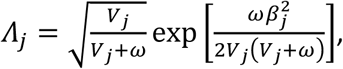

where β*_j_* and *V_j_* denote the estimated allelic effect (log-OR) and corresponding variance from the meta-analysis. The parameter *ω* denotes the prior variance in allelic effects, taken here to be 0.04^57^.

We then calculated the posterior probability that the *j*th variant drives the association signal, given by

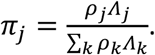

In this expression, *ρ_j_* denotes the prior probability that the *j*th variant drives the association signal, and the summation in the denominator is over all variants across the locus. We considered two prior models: (i) functionally unweighted, for which *ρ_j_* = 1 for all variants; and (ii) annotation informed, for which *ρ_j_* is determined by the functional severity of the variant. For the annotation informed prior, we considered five categories of variation^16^, such that: (i) *ρ_j_* = 165 for PTVs; (ii) *ρ_j_* = 33 for moderate-impact variants; (iii) *ρ_j_* = 3 for low-impact variants; (iv) *ρ_j_* = 1.5 for other variants mapping to DHS; and (v) *ρ_j_* = 0.5 for all other variants.

For each locus, the 99% credible set^22^ under each prior was then constructed by: (i) ranking all variants according to their posterior probability of driving the association signal; and (ii) including ranked variants until their cumulative posterior probability of driving the association attained or exceeded 0.99.

### Functional impact of coding alleles

We used CADD^37^ to obtain scaled Combined Annotation Dependent Depletion score (CADD-score) for each of the 40 significantly associated coding variants. The CADD method objectively integrates a range of different annotation metrics into a single measure (CADD-score), providing an estimate of deleteriousness for all known variants and an overall rank for this metric across the genome. We obtained the estimates of the intolerance of a gene to harbouring loss-of-function variants (pLI) from the ExAC data set^36^. We used the Kolmogorov-Smirnov test to determine whether fine-mapping groups 1 and 2 have the same statistical distribution for each of these parameters.

### T2D loci and physiological classification

To explore the different patterns of association between T2D and other anthropometric/metabolic/endocrine traits and diseases, we performed hierarchical clustering analysis. We obtained association summary statistics for a range of metabolic traits and other outcomes for 94 coding and non-coding variants that were significantly associated with T2D through collaboration or by querying publically available GWAS meta-analysis datasets. The z-score (allelic effect/SE) was aligned to the T2D-risk allele. We obtained the distance matrix amongst z-score of the loci/traits using the Euclidean measure and performed clustering using the complete agglomeration method. Clustering was visualised by constructing a dendogram and heatmap

### URLs

Type 2 Diabetes Knowledge Portal: http://www.type2diabetesgenetics.org/

